# Spatial and semantic regularities produce interactive effects in early stages of visual orientation

**DOI:** 10.1101/639385

**Authors:** Giuseppe Notaro, Uri Hasson

**Affiliations:** Center for Mind/Brain Sciences (CIMeC), The University of Trento, Italy

## Abstract

Learning environmental regularities allows predicting multiple dimensions of future events such as their location and semantic features. However, few studies have examined how multi-dimensional predictions are implemented, and mechanistic accounts are absent. Using eye tracking study, we evaluated whether predictions of object-location and object-category interact during the earliest stages of orientation. We presented stochastic series so that across four conditions, participants could predict either the location of the next image, its semantic category, both dimensions, or neither. Participants observed images in absence of any task. We modeled saccade latencies using ELATER, a rise-to-threshold model that accounts for accumulation rate (AR), variance of AR over trials, and decision threshold. The main findings were: 1) accumulation-rate scaled with the degree of surprise associated with location of target-presentation (confirmatory result); 2) predictability of semantic-category hindered latencies, but only when images were presented at a surprising location, suggesting a bottleneck in implementing joint predictions; 3) saccades to images that satisfied semantic expectations were associated with larger variance of accumulation-rate than saccades to semantically-surprising images, consistent with a richer repertoire of early evaluative processes for semantically-expected images. Joint impacts of location and target-identity regularity were also identified in analyses of anticipatory fixation offsets. The results indicate a strong interaction between the processing of regularities in object location and identity during stimulus-guided saccades, and suggest these regularities also impact anticipatory, non-stimulus-guided processes.

## Introduction

### The question

The ability to learn about the environment and adapt behavior accordingly is a fundamental human capacity. This knowledge can support predictions about the future, and these predictions are inherently multidimensional. While walking in a crowded space, biking through dense traffic, or playing a video game, individuals can anticipate where future events will take place (e.g., traffic will be coming from right), what attributes these events will have (it’s likely to be a large garbage truck), as well as other dimensions such as the timing and duration of the expected events.

Prior knowledge about the location of future events facilitates processing. In experimental studies, people respond faster to targets that appear in more likely locations (e.g., Geng & Behrmann, 2002). Foreknowledge of a target’s location increases firing rates for visual neurons whose receptive fields overlap with the expected location, which is further associated with faster saccades towards the target (e.g., Basso & Wurtz, 1998). In humans, statistically-derived probabilistic information about a future target’s location has recently been shown to produce anticipatory fixation offsets prior to target appearance, which account for considerable variance in the subsequent saccade latencies to target (Notaro, van Zoest, Altman, Melcher, & Hasson, 2019). Less is known about how foreknowledge of a target’s semantic/perceptual features (*identity* henceforth) impacts processing, particularly with respect to how identity expectations impact saccades to target. Expectations about shape (Deubel & Schneider, 1996), shape-and-color (Kingstone, 2013) or motion (Tanaka, Yoshida, & Fukushima, 1998) also impact saccade latency. Related work has shown that providing definitive information about a future target’s identity facilitates its identification, as seen in an increased proportion of correct first gazes to such targets when embedded within distractors (for examples of such category-driven visual search effects, see Moores, Laiti, & Chelazzi, 2003; Yang & Zelinsky, 2009).

What is not known is how and whether multidimensional predictions – ones that pertain to both location and identity – are implemented, and whether those at all optimize behavior. In many circumstances, the location and identity of future events can be probabilistically anticipated. However, we do not know how such knowledge impacts orientation to targets, and as a result, even core questions such as whether the ability to predict both location and identity facilitates or hinders processing, remain unaddressed. What *is* known, as we detail below, pertains a different issue; namely, how people make decisions when required to engage in deliberative decisions about target-features. This line of work studies decision-making processes that: are inherently late-occurring; *ii*) have been linked to other functional systems (e.g., Rajan et al., 2018); and *iii*) may not constitute an integral component of natural perception.

To study how multidimensional predictions impact behavior, we designed an eye-tracking study where statistical features of the environment, manipulated via a first-order Markov process, could produce independent probabilistic expectations about the location or identity of the next visual target that would appear. In brief, we constructed different stimulus-series so that some series carried statistical information about either the next target’s location; its identity; its location and identity; or neither location nor identity. This allowed us to determine whether the existence of regularities in identity or location (what/where) produce signatures of non-independence during early stages of stimulus processing.

We concentrated on saccade latencies because these are among the earliest observable stages of behavioral orientation. As detailed below, while learning of statistical regularities (statistical learning) is thought to give rise to predictions (e.g., Hasson, 2017; Notaro et al., 2019), it is unclear how these predictions impact different stages of processing. As mentioned, prior studies required explicit decisions, usually communicated via key-press. This effectively integrates over early and late sensory processing as well as executive processing stages and cannot isolate the impact of probabilistic foreknowledge on early orientation. Furthermore, as outlined below, the modest extant literature on this topic has produced conflicting conclusions on whether learning location/identity sequences (or using location/identity information for prediction) relies on independent or interactive systems.

### Theoretical context and related work

Little is known about how individuals learn statistics of an environment when those statistics independently pertain to the location and identity of future events. Some studies suggest there exist separate functional systems for processing regularities that govern target location and identity. These studies used the Serial Reaction Time (SRT) paradigm to study how effectively individuals learn regularities that govern target locations, in conditions where the identity of the presented targets (which constituted an irrelevant dimension) followed either a regular or random pattern (deterministic sequences Mayr, 1996; Deroost & Soetens, 2006; probabilistic sequences: Remillard, 2017). Those studies showed that that participants learned the structure of both target-location and target-identity sequences. Importantly, that there was no penalty for learning location-regularity when the sequence of target identities was regular as opposed to when it was random. This suggests that learning the identity stream did not produce a processing cost. A strength of these studies is that knowledge of location and identity is attained endogenously rather than cued via explicit prompts, which improves external validity with respect to studying knowledge attainment and use.

Others have used cue-target paradigms to determine whether providing explicit cues about identity and/or location of a future target produces additive or interactive effects. Mattler (2003) cued participants to the location or identity of a future target and found that valid cues (producing expected trials), for both location and identity, resulted in faster responses than invalid cues (producing surprisal trials). These effects were non-additive: the cue-validity effect for target-identity (valid – non valid cue regarding identity) was stronger when location cues were valid than when location cues were invalid. This was interpreted as indicating the identity-related feature maps exert an early effect that interacts with spatial attention processes. Bruhn and Bundesen (2012) used a similar design, but arrived at a different conclusion. ^1^ In three experiments, they consistently identified beneficial effects for both location and identity cuing but no interactive effects. They concluded that this suggests serial stages of processing that are functionally independent (see Remillard, 2017, for similar proposal). Egner et al. (2008) used a similar design but also collected neuroimaging data, and both the behavioral and neuroimaging data suggested largely additive effects of location and identity cuing.

Taken together these behavioral studies are consistent with the idea that location and identity information are processed almost independently, at least for the reported experimental paradigms. Such findings appear consistent, *prima facie*, with the canonical dual-stream neurobiological approach to the visual system (Ungerleider & Haxby, 1994). However, later neuroimaging studies suggest a more nuanced and interactive view of location and identity processing (e.g., Cichy, Chen, & Haynes, 2011; Davis & Hasson, 2018).

### Current approach

Given the above-reviewed state of the art, we were interested in how endogenously derived representations of statistical regularities, separately pertaining to location and identity, impact saccade latencies to targets. Crucially, in our study participants were requested to saccade toward the presented images without making any decision about the images’ identity. For this reason, from the perspective of perceptual decisions, the saccade latencies (SL) only reflected a decision about the gaze target (Carpenter & Williams, 1995). Consequently, any eventual modulation of SL due to the identity regularities would be interpreted as an automatic, non-demand-driven interaction between functional systems involved in planning gaze location and those involved in implementing predictions regarding target identity. We used a computational decision-accumulation model (ELATER; Nakahara, Nakamura, & Hikosaka, 2006) to model the distribution of SL saccade latencies. This is a simple model capable of separating two stages of processing; the first pertaining to the accumulation of sensory evidence, and the second to the decision *per se*. ELATER estimates the following independent parameters: accumulation rate, the variance of the accumulation rate, and the variance of the decision threshold.

Our first analysis aimed to demonstrate the feasibility of ELATER modeling and evaluated whether presentation of targets in high-probability locations produced higher accumulation rates. This would confirm canonical results in this area. Our second and third analyses evaluated two questions:

1. In series with regular identity transitions, whether expected and surprising identity-transitions had different impact on saccade latencies. We term this a *specific* effect of identity regularity as it differentiates expected-identity from surprising-identity trials and thus shows sensitivity to the identity of a target.
2. Whether having knowledge about the identity of a future target could in itself produce processing costs as compared to conditions that did not allow an identity-prediction. We term this textcolorbluecontrast between regular and regular identity series, a *non-specific* effect of identity as it shows that individuals have identified the regularity in the identity-stream but without using this knowledge on a trial-by-trial basis.

## Methods

### Participants

Forty volunteers participated in the study (mean *Age* = 23.02 ±.75; SEM is the measure of spread throughout unless noted otherwise). They were recruited from the local student community and reimbursed 15 Euro for their time. The Ethical Review Board approved the study.

### Design

The design of the study consisted of two factors: one varying the expectation of the target location and one varying the expectation of the target identity. For the Location factor this meant that in one case the target was presented in a location that was statistically-expected (high-frequency transition), in the second case the target was presented in a surprising location that violated expectation (low-frequency transition), and in the third case it was presented in a series where all location transitions were random (we call this a nonRegular (not-predictable) location trial. The same held for the Identity factor: in one case the target identity violated statistical expectation (surprising), in other case it satisfied expectation (expected), and in the other case it was nonRegular. The principle of the design was to cross these two factors in order to generate nine conditions that independently varied in the surprisal of location and identity. As we detail below, the design also allows separating typical signatures of surprisal (*surprising* ≠ *expected*) from a more general and less specific signature of learning where the surprising and expected trials do not dissociate, yet both differ from nonRegular trials: ([surprising ≈ expected] ≠ nonRegular)

To produce these nine conditions, we created four types of series (see also Figure 1A):

**Figure 1.**
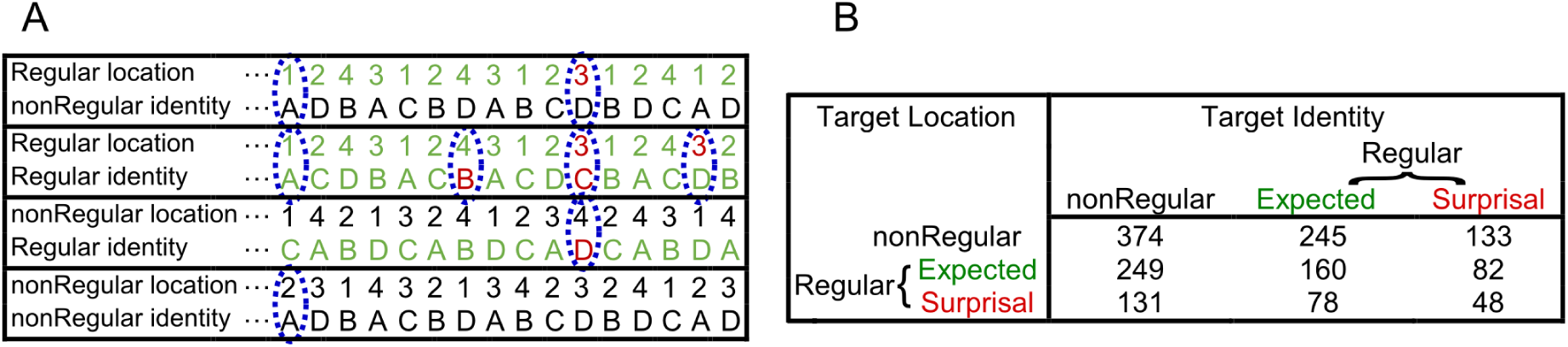
Study design. **Panel A.** Examples of the four types of series used in the study. These were created by crossing the predictability of Location and Identity transitions within each series. Series with non-regular (nonRegular) transitions are presented in black. For series with regular transitions, expected trials are presented in green and surprising trials are presented in red. The blue ellipses highlight the nine types of experimental trials. **Panel B.** Mean number of trials, part participant, for each of the nine types of experimental trials.

1. Series where there was no regularity in either the location transitions or identity transitions (nonRegular Location / nonRegular Identity in Figure 1A).
2. Series where only location transitions were regular. This produced expected location transitions and surprising location transitions, but all identity transitions were random (Regular Location / NonRegular Identity in Figure 1A).
3. Series where only identity transitions were regular. This produced expected identity transitions and surprising identity transitions, but all location transitions were random (nonRegular Location / Regular Identity in Figure 1A).
4. Series where both location and identity transitions were regular. This produced four types of trials within these series, so that: *i*) both location and identity transition were expected; *ii*) location transition was expected and the identity transition was surprising; *iii*) location transition was surprising and the identity transition was expected; *iv*) both the location and identity transitions were surprising (Regular Location / Regular Identity in Figure 1A.)

For each of these four types of series, participants were presented with four series, each consisting of 128 trials. To limit the effect of the inhibition of return we did not allow repetitions of location or identity across successive trials by setting the transition matrices so that the diagonal elements were set to zero. To generate the non regular transitions we used the transition matrix *M*_*nonReg*_ with equi-probability for all transitions (*p* = 0.33) (see below). To generate the regular transitions we used transitions with *p* = 0.66 and *p* = 0.33 probabilities. We created four different types of Regular matrices (see example below), so that when averaging across the different types of spatial pair-wise transitions, the mean transition matrix of all these Regular matrices was identical to that of the nonRegular condition. This means there was no confounding between the proportion of specific types of saccades and the experimental condition (see Appendix for further details).

Example of a Regular (*M*_*Reg*1_) and the nonRegular transition matrices (*M*_*nonReg*_) are respectively:

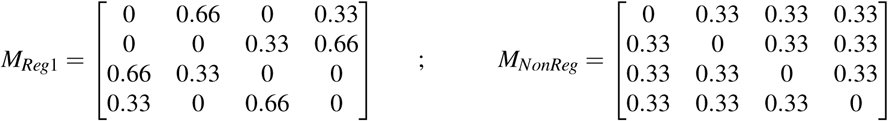

Markov entropy was 0.92 bits/trial for the *Regular* process and 1.58 bits/trial for the *nonReg* process. Note that, as described above, such transition matrices independently determined transitions between spatial locations or image-categories, as specified in the different conditions, so that the statistical features of the identity and location streams needed to be tracked separately. From these transition matrices we produced series with 120 trials. In the study, these series were presented according to a random order determined separately per participant. At the end of each 120-trial series we added 8 trials with random images presented in clockwise or anti-clockwise manner to partially reduce the impact of recent statistical structure. In addition, the first 8 trials of each series were not analyzed as by definition the *Regular* and *nonRegular* series cannot be discriminated immediately.

The four types of series described above were constructed to generate the 9 types trials types in Figure 1B. Beyond this, they were not used as factors in the design or referred to in contexts of statistical analyses. To evaluate statistical interactions between identity and location knowledge, we analyzed the 9 trial-types in the analysis matrix presented Figure 1B, using 3 × 3 interactions and follow-up analyses. In Figure 1B, each cell presents the mean number of trials, *per person*, for each cell in the experimental design.

## Procedure

### Eye-tracking

Stimuli were displayed on a CRT display (Diamond pro 2070SB, Mitsubishi Electric Corporation, Tokyo, Japan) with a spatial resolution of 1280 × 1024 pixels, and a 75 Hz refresh rate. We generated the experimental software using Matlab™ and the Psychophysics Toolbox extensions (Brainard, 1997). Participants’ eyes were set at the same height as the screen center and at a distance of 58 cm. Eye position signals were recorded by a separate computer with a tower-mounted, video-based eye tracker (*Eyelink 1000* Tower mount, SR Research Ltd, Mississauga, Canada) and were sampled monocularly at 1000 Hz. We performed a nine-point calibration procedure during which the eye-tracker calculated a mapping between sensor and display positions.We performed calibration after each break. Before beginning the experiment we identified each participant’s dominant eye using the Dolman method.

### Stimuli and trial structure

The timeline of each trial was as follows (see Figure 2A): a fixation symbol appeared for 400 ms, followed by a post-fixation blank screen for 160 ms. A target was then presented for 360 ms, and and a post-target blank screen for 160 ms. The fixation symbol consisted of an inner gray circle with a radius of 0.4° (same color as background) within an outer black circle with a radius of 1.2°. We chose this fixation symbol as it has been shown to allow some variance in eye movement during fixation (Thaler, Schütz, Goodale, & Gegenfurtner, 2013).

**Figure 2.**
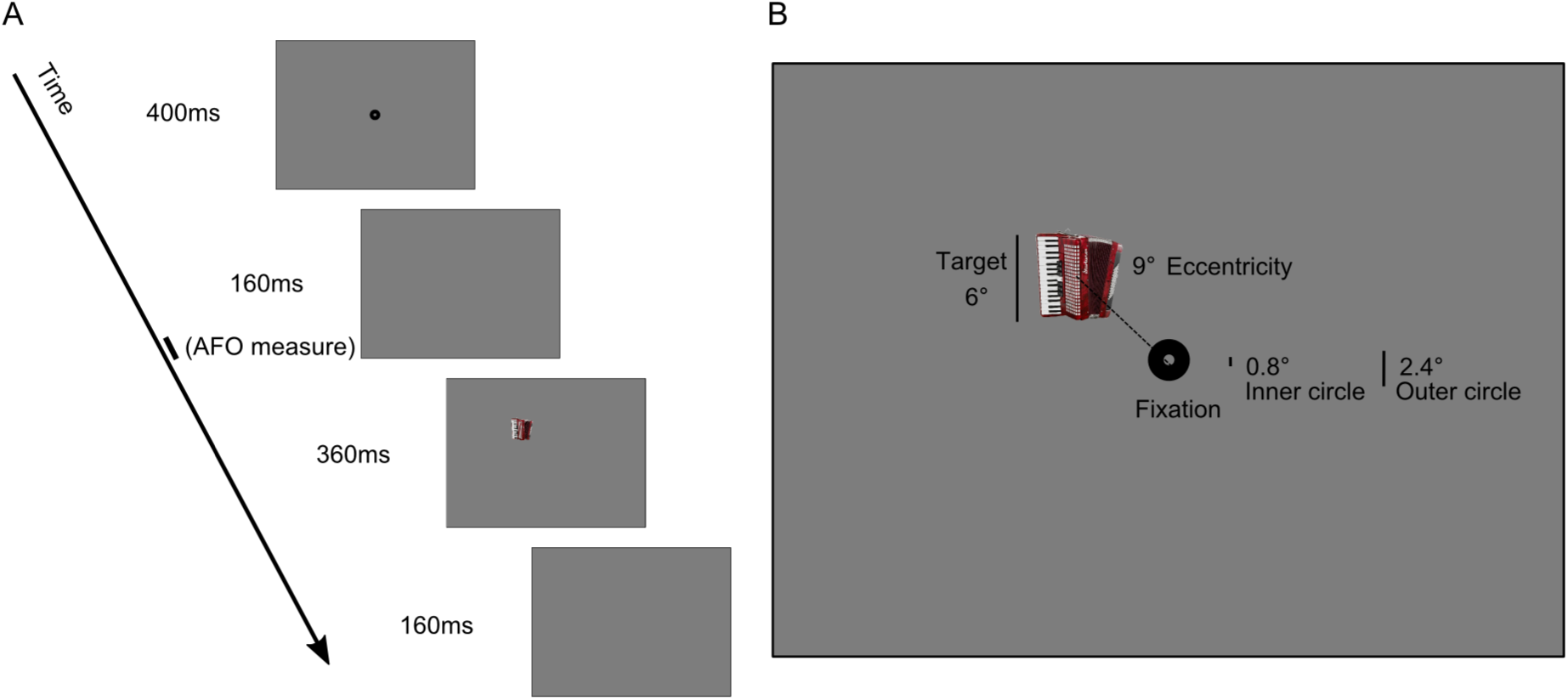
Experimental setup.**Panel A**: Timeline. **Panel B**: Location of display elements.

The visual targets appeared at one of four screen quadrants (top left, top right, lower left, lower right), at an eccentricity of 9°from screen center. Targets subtended a visual angle of 6.36 degrees in each screen coordinate. Participants were instructed to saccade rapidly to the target and fixation symbol when they appeared (see Figure 2).

The target images belonged to one of four categories of RGB images with 125 different elements each: neutral faces, musical instruments, fruits and tools. The faces were taken from the Center for Vital Longevity Face Database (Minear & Park, 2004) and from SiblingsDB, a face database for machine learning studies (Vieira, Bottino, Laurentini, & De Simone, 2014). All the other images were retrieved from publicly accessible online repositories. When assigning these images to the 120-trial series, the elements were assigned randomly with the additional following constraints: the same image could be presented only once per series, could not be presented in two consecutive series, and could not be presented in the same location as the previous time it was presented. A subset of 5 images was used only in the last 8 trials of each session where the location was deterministic. A fifth category of images (‘Letters and numbers’) was used during the training before the beginning of the experiment (see below).

### Instructions and training

Participants were instructed to maintain their gaze on the fixation symbol when it was present and to move their gaze quickly toward the presented image. The instructions did not mention image-identity. To maintain participants’ alertness, we included catch trials that consisted of black discs with a white line through them that appeared instead of the regular fixation symbol. Catch trials appeared every 23–27 trials following a uniform distribution. Participants were told that catch trials would appear infrequently and that they were to press the mouse button when they saw them. Following each series, participants were presented with performance indicators for that series, which included the number of targets and fixation symbols saccaded to within an allowed spatial and temporal tolerance (see below), the number of correct catch trials and eye blinks, as well as their overall mean performance to that point. This was done to motivate participants to perform well and to provide a buffer between the stochastic contexts of the just completed and the following series.

Before beginning the study proper, participants underwent training where they viewed series of 20 trials each, until they were comfortable with the procedure (typically within 2–7 series). The training session differed in some respects from the main experiment. The images were of the same size as those in the main study, but contained a monochromatic letter or number, to avoid exposing participants to the same image-categories used in the experiment. In the training series there were no transition constraints (each transition had probability of 25%) so that participants could not develop experience with the transition structure used in the study proper. In addition, during training (but not the study proper) we provided real-time positive auditory feedback when participants’ gaze hit the target or the fixation circle within 200 ms from appearance and with a maximum deviation of 1°from their borders, and whenever participants correctly responded to catch trials with a mouse click. We provided negative auditory feedback whenever participants failed to hit the target, failed to respond to catch trials, or blinked. While participants were instructed and trained to arrive at fixation within 200ms from appearance, we did not penalize participants for saccading to the screen center prior to stimulus appearance. Here and in the rest of the text, the term “anticipatory saccades” refers to saccades to the laterilized targets (which were of main interest) rather than saccades to central fixation. A summary of the positive and negative scores was presented at the end of each training session.

### Analysis

#### Saccade classification

We detected saccades by adaptively determining speed thresholds relative to saccade onset, offset and peak (Nyström & Holmqvist, 2010). We defined saccade onset (offset) time as the time of the first local minimum with speed below an adaptive threshold, preceding (following) a saccade peak.

#### Anticipatory fixation offsets and trial selection

We defined a measure of gaze location on each trial (Gaze Offset) as the mean gaze in the *x* direction in the last 10 milliseconds prior to target appearance. These Gaze Offset data were the input to the steady-state analysis of gaze location described below.

In a subsequent analyses, for convenience, we coded gaze offset as positive if in the same direction of the previous target x-coordinate, as negative otherwise. We refer to this quantity as Anticipatory Fixation Offset (AFO). Since AFOs are small in amplitude (usually < 0.5°), to avoid having a few outliers bias the mean AFO value, we analyzed only trials in which AFO was below 3°(as in Notaro et al., 2019). Other than this constraint, we also analyzed only these trials in which the measured saccades were reactive to the target presentation. This was defined as having a latency above 80 ms (see Fischer & Boch, 1983) and below target offset (400 ms) and that saccades landed close to the image center (Melcher & Kowler, 1999) with an allowed error of 3.5°. This was done in order to include only stimulus-guided saccades and exclude anticipatory ones. Finally, we excluded the first eight trials of each series, since the stochastic content could not be discriminated immediately. Given these constraints, the average percentage of valid trials was 83.3 ± 2.1%.

#### Steady state analysis of gaze offsets

The series with regular and non-regular location-transitions have different recurrence patterns because they are generated by different Markov processes. As detailed in the *Appendix*, for the nonRegular process the modal recurrence time is 2 trials because the most probable event is to alternate screen side on each trial as two of the three potential targets are on the alternate side. In contrast, for the Regular series the modal recurrence is 3. This difference in recurrence directly translates into different power spectra of target location, see Figure Appx.1A. If these recurrence patterns drive anticipatory gaze, this will produce different power spectra when the anticipatory gaze positions (one datum per trial) are analyzed in the frequency domain.

The steady-state analysis is essentially model-free which means that it can identify signatures of regularity in the time series of eye data even in absence of a specific *a-priori* model of how learning and predictions occur as result of complex computations that cannot be hypothesized in advance. Such complex integration/prediction patterns potentially can annul any direct relation between anticipatory gaze and the most probable next location. As discussed in the *Appendix*, section: *Gaze offsets does not directly signal screen side of most likely subsequent target*, a simple ANOVA, which evaluated whether the gaze tracked the position of the next high-probability target location, did not identify such a direct relation.

As input to the power spectra analysis we considered, separately, the horizontal (*X)* and vertical (*Y)* gaze coordinates of the single Gaze Offset measurement obtained in each trial, that is, the mean gaze position 10ms prior to target appearance.^2^ To accomodate missing Gaze Offset values in non–valid trials we used the following procedure:

1. We obtained a unique time series (*x*) for each participant and condition by concatenating the single gaze-offset measures per trial and separating each two sessions by a series of 12 *nan* values.
2. We calculated the autocorrelation function up to the maximum lag of 11 trials: *R*_*xx*_ (*τ*) = ∑_*k,*(*k*−*τ*)*∈*[*valid trials*]_ *x* (*k*) *x* (*k* −*τ*), where *τ* = 0, *…* 11.
3. We calculated the power spectral density *S*_*x*_ as the Fourier transform of *R*_*xx*_ (Wiener–Khinchin theorem, see Engelberg, 2006).
4. We evaluated the Frequency Tagged Response (*FTR*_*x*_) at *f*_*tag*_ = 1*/*3 *trials*^−1^, per participant. This was operationalized as the power density at *f*_*tag*_ divided by the mean power in the four closest frequency bins.

#### Implementation of ELATER

The LATER model (Carpenter & Williams, 1995) is based on the experimental finding that while saccade latency (SL) is a variable that has a skewed distribution, the distribution of its reciprocal (promptness) is symmetric and well described by a Gaussian. Consequently, the reciprocal has two free parameters, *µ* and σ_*µ*_ describing its normal distribution. In LATER, these parameters are taken to describe a generative decision process, where the sensory evidence is accumulated with a rate *µ*, constant within a trial, but variable across trials according to the uncertainty σ_*µ*_. A decision is taken when the accumulated evidence reaches a threshold, generally set to 1.

LATER predicts that SLs lies on a straight line in a reci-probit plot and the model parameters determine the line’s intercept (*µ*/σ_*µ*_) and slope (1/σ_*µ*_). In some cases the model fails to include the lower values of the distribution, which are treated as *early responses*. Early responses lie on a separate line when plotted on the reci-probit plot and are typically ignored when determining the main parameters (Appx.2B, dashed blue line). However, trimming out these responses could produce a loss of information otherwise available in the entire SL distribution (Heathcote, Popiel, & Mewhort, 1991).

The ELATER model (Nakahara et al., 2006) extends LATER with an additional free-parameter: the variability of the starting point of the decision process (*sigma*_0_), see Figure Appx.2A. When *sigma*_0_ is fixed at zero, this model reduces to LATER, and SLs lie on a straight line in a reci-probit plot. But for higher values of *sigma*_0_, the line bends where SL values are smaller, and the resulting curve therefore includes early responses as well (see Figure Appx.2B, red line).

We fit ELATER model parameters for each participant, for each of the 9 cells determined by the experimental condition. To estimate the model parameters we used a global search algorithm (function *fminsearch* in Matlab) that minimized the errors in quantiles of probabilities, spaced at 0.05. As the starting point for the algorithm we used the solution from the LATER model, obtained by performing a linear fit in the reci-probit plot and setting *sigma*_0_ = 0. In several follow-up analyses we evaluated non-specific effects of identity regularity. In those cases, we considered the expected and surprising conditions as a single condition and fit a single parameter to to the resulting distribution. The advantage of this procedure over averaging parameter estimates for expected and surprising trials is that it considers more trials in the parameter estimation and is theoretically motivated^3^.

Before conducting analyses of variance we tested for homoscedasticity of variance across the nine design cells, for each of the three model parameters. Bartlett’s test indicated a rejection of homoscedasticity for each of the three parameters (all *p* < .001). For this reason we applied a non-parametric test which applies the analysis of variance to the aligned-rank-transformed data (ART, Wobbrock, Findlater, Gergle, and Higgins, 2011). We also verified whether the statistical conclusions held without the transformation and no exceptions were found. In each of our analysis, we conducted a 3 (Location: nonRegular, expected, surprising) × 3 (Identity: nonRegular, expected, surprising) ANOVA to identify main effects and interactions. We then conducted follow-up analyses, Bonferroni corrected for multiple comparisons, that probed for specific and non-specific effects of series-regularity.

## Results

### Saccade latencies

A 3 (Location: nonRegular, expected, surprising) × 3 (Identity: nonRegular, expected, surprising) ANOVA of mean SL showed a main effect of Location, *F*(2, 39) = 50.12, *p* < .001. Figure 3A reports the SL values for all experimental cells. As shown in the Figure, SL increased with the degree of location-surprise. Namely, SL were lowest for expected transitions (*M* = 144.76 ± 1.75*ms*), higher for nonRegular transitions (*M* = 147.76 ± 1.82) and highest for surprising transitions (*M* = 150.07 ± 1.90). Collapsing over Identity, pairwise T-tests demonstrated differences between expected-location and nonRegular-location trials (*t*(39) = 5.47, *p* < .001, *d* = 0.27) as well as between nonRegular-location and surprising-location trials (*t*(39) = 3.65, *p* < .001, *d* = 0.20). We note that SL was quite rapid given that only a single target was presented on the screen, without distractors, and with a sufficient gap interval between the fixation and saccade.

**Figure 3.**
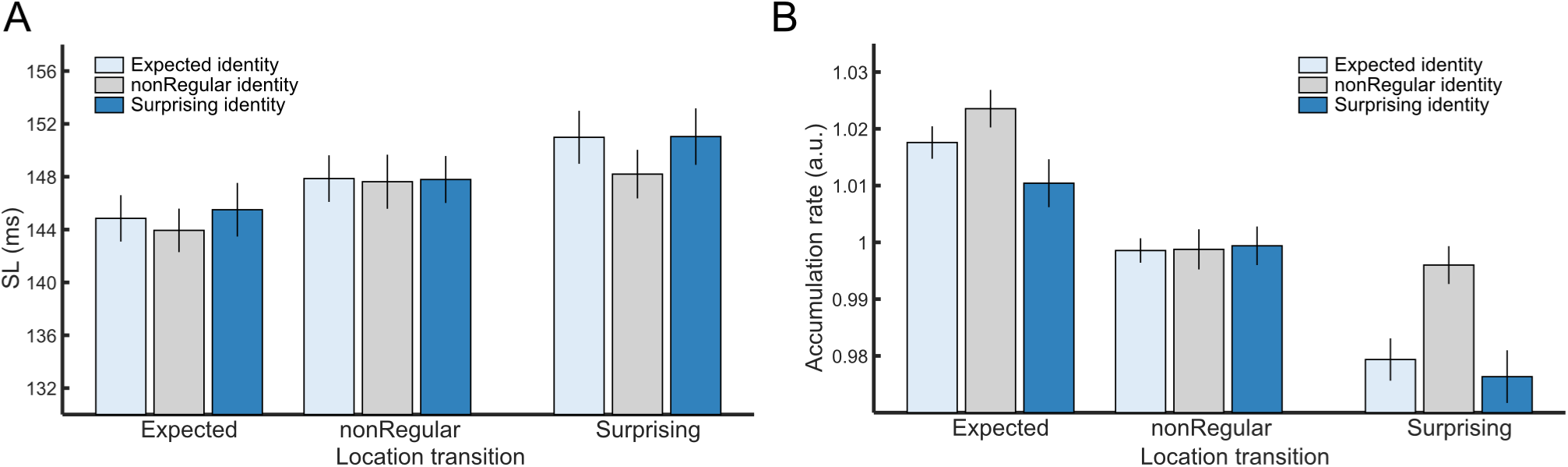
Mean saccade latencies and ELATER-estimated accumulation rates in the nine experimental conditions. **Panel A.** Mean saccade latencies. The analysis of saccade latencies identified a main effect of Location and a main effect of Identity. **Panel B.** Mean accumulation rate. The analyses of accumulation rate revealed main effects of Location, Identity and an interaction between the factors.

There was also a main effect of Identity, *F*(2, 39) = 4.82, *p* = 0.009 (latencies were *M* = 147.90 ± 1.78*ms* for expected trials; 146.59 ± 1.80*ms* for nonRegular trials, and 148.11 ± 1.88*ms* for surprising trials.) This main effect notwithstanding, the pattern of SL did not track levels of surprisal. The two-way interaction between Location and Identity did not approach significance, *F*(4, 39) = 1.45, *p* = .22.

### ELATER model of saccade latencies

#### Accumulation rate

In ELATER, accumulation rate (*µ*) describes the rate by which external information about a target is accumulated. Mean *µ* across participants closely tracked the pattern found for SL, and is depicted in Figure 3B. A 3 (Location) × 3 (Identity) ANOVA identified a statistically significant effect of Location (*F*(2, 39) = 85.13, *p* < .001). Collapsing over Identity, we found that *µ* tracked location surprisal. It was largest (reflecting faster accumulation) when location was expected, lower when location was nonRegular, and lowest when location was surprising (post-hoc comparisons: expected vs. nonRegular, *t*(39) = 6.11, *p* < .001, *d* = 1.70; nonRegular vs. surpising, *t*(39) = 4.87, *p* < .001, *d* = 1.27), see Figure 4A.

**Figure 4.**
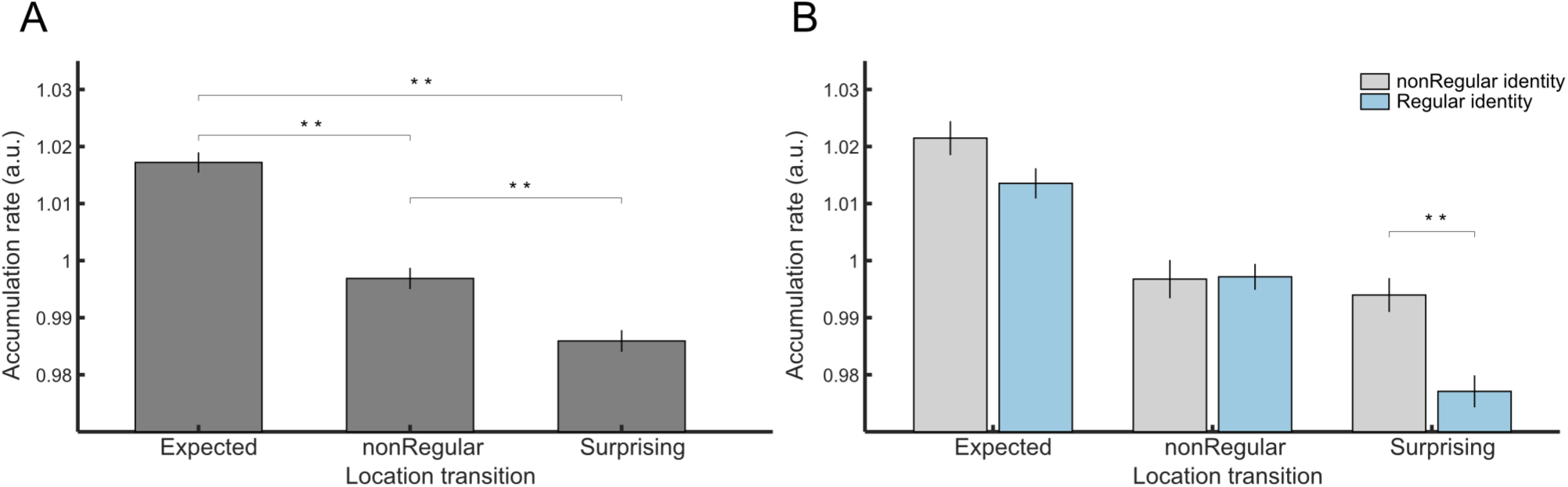
ELATER-estimated accumulation rates. **Panel A.** Mean accumulation rate for saccades to expected, surprising, and nonRegular locations (data are collapsed across levels of the Identity factor). **Panel B.** Mean accumulation rate for expected, surprising or nonRegular locations as function of whether identities were presented within statistically-regular or non-regular series. Two asterisks indicate significant pairwise differences with *p* < .001 (Bonferroni corrected).

There was also a significant effect of Identity, *F*(2, 39) = 10.47, *p* < .001. Collapsing across Location, we found that *µ* was highest in the nonRegular condition, and significantly higher than both the expected (*t*(39) = 3.36, *p* = .0035, *d* = 0.86) and surprising conditions (*t*(39) = 3.79, *p* < .001, *d* = 1.10); see Figure 4B. There was however no difference between the identity-expected and identity-surprising trials, even when examined separately within each level of location predictability (all *ps* > .1). The effect of identity therefore does not track identity-surprisal but is instead consistent with a non-specific effect of identity regularity, as identified for the analysis of saccade latencies. To summarize, to this point, the data for accumulation rate match those found for the mean saccade latencies.

Departing from the findings reported for SL, accumulation rates reflected a significant interaction between the two factors. *F*(4, 39) = 2.95, *p* = 0.02. As shown in Figure 4B, presentation of targets in series where identity was regular (i.e., collapsing over surprising and expected trials) negatively impacted *µ* (indicating less effective accumulation), but only when target-location was surprising. This stood in marked contrast to when targets were presented at nonRegular locations, in which case being able to predict target identity had almost no effect.

To verify these patterns statistically, we defined an identity-regularity cost (*idcost*) as the difference between the values of *µ* for the nonRegular identity trials on the one hand, and the combined set of expected and surprising identity trials on the other. We then computed this cost within each level of location trials (nonRegular: *idcost*_nonReg_; expected: *idcost*_exp_; surprising: *idcost*_surp_). Consistent with the interaction term, *idcost*_surp_ was statistically significant, *M* = −0.0169 ± 0.004*a.u., t*(39) = 3.91, *p* < .001, *d* = 0.62, but *idcost*_exp_ was not (*M* = −0.008 ± 0.004*a.u., t*(39) = 1.86, *p* = .070, *d* = 0.29), and *idcost*_nonReg_ was not either (*M* = 0.0004 ± 0.004*a.u., t*(39) = 1.10, *p* > .1, *d* = 0.016). Figure 4B presents these pair-wise comparisons. As we detail in the Discussion, this suggests a general bottleneck in processing surprising locations when identity predictions are licensed.

Though *idcost*_exp_ was not significant, the magnitude of *idcost*_exp_ correlated (across participants) with *idcost*_surp_, (*rho* = 0.41, *p* = .0079), as shown in Figure 5. Conversely, *idcost*_surp_ did not correlate, across participants, with *idcost*_nonReg_ (*idcost*_exp_ did not correlate with *idcost*_nonReg_ either) (*ps* > .1). This suggests that a common factor may determine the impact of identity-regularity when orienting to both surprising and expected locations.^4^

**Figure 5.**
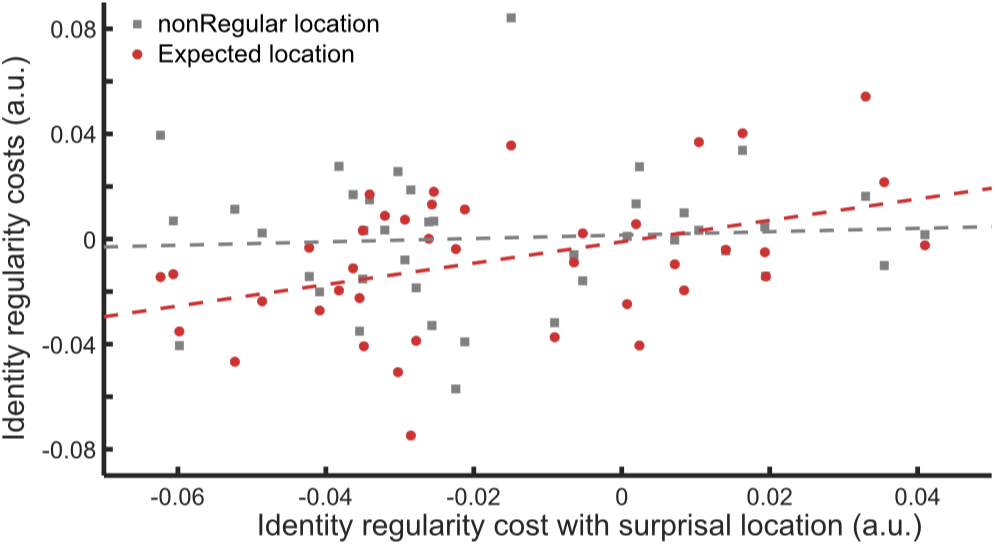
Scatterplots of identity costs: nonRegular vs. surprising locations (grey squares, grey best fit line); expected vs. surprising locations (red circles, red best fit line). Identity costs are calculated as differences in accumulation rates and are in arbitrary units. We found similar correlations when defining identity costs as differences in saccade latencies.

#### Evaluating identity-regularity costs against saccade latencies

Slower saccade latencies are known to more strongly reflect top-down modulations than faster latencies (e.g., Van Zoest & Donk, 2008; Schütz, Trommershäuser, & Gegenfurtner, 2012). We found identity-regularity costs for saccades made to surprising locations, but these saccades were also slower than saccades to expected or non-predictable locations. Therefore, to determine whether the crucial factor for the impact of identity-regularity was location-suprisal or saccade-latency, we performed for each participant median splits of the saccade latency distribution within each level of location (surprising, expected, or nonRegular). We then evaluated if the identity-regularity cost was larger for the slower than faster saccades: for each condition we calculated the identity-regularity cost in the upper and lower sets constructed by the median split, here defined as the SL latency-difference between regular and non-regular identity transitions.^5^

When location was surprising, identity-costs were positive, but neither the upper nor lower splits departed from zero (slower: *M* = 1.1 ± 1.0 ms, *p* > 0.1; faster: *M* = 0.60 ±.46 ms, *p* > 0.1) and they did not significantly differ (*p* > 0.1). A similar pattern held when location was expected (slower: *M* = 0.69 ± 0.69 ms, *p* > 0.1; faster: *M* = 0.62 ±.31 ms, *t*(39) = 2.02, *p* = .051, *d* = 0.32; no significant difference). When location was nonRegular, identity costs for the slower-saccade split were positive, though not significantly so (*M* = 1.20 ±.63 ms, *t*(39) = 1.89, *p* = .066, *d* = 0.30), whereas for the faster split identity costs were negative, indicating faster saccades for regular-identity transitions (*M* = − 0.62 ±.27 ms, *t*(39) = 2.31, *p* = .026, *d* = 0.37). This produced a significant difference between identity costs for slow and fast saccades in the nonRegular condition, *t*(39) = 2.84, *p* = .007, *d* = 0.45. These findings do not support with the notion that identity-regularity costs (penalties) are more strongly expressed for slower saccades, and suggest there are some cases where these regularities can speed up processing (as seen for faster saccades in the non-regular location condition).

#### Variance of Accumulation Rate

We implemented a 3 (Location) × 3 (Identity) ANOVA to analyze how these factors impacted the variance of the accumulation rate (σ_*µ*_), considering the estimated values for each participant, in each of the nine conditions (Fig. 6A). The main effect of Location was not significant (*p* > .1). The main effect of Identity was significant, *F*(2, 39) = 4.33, *p* = .014. Post-hoc comparisons showed this was due to a significantly higher σ_*µ*_ values for expected trials than surprising trials, *t*(39) = 3.08, *p* = .023, *d* = 0.77, see Figure 6A

**Figure 6.**
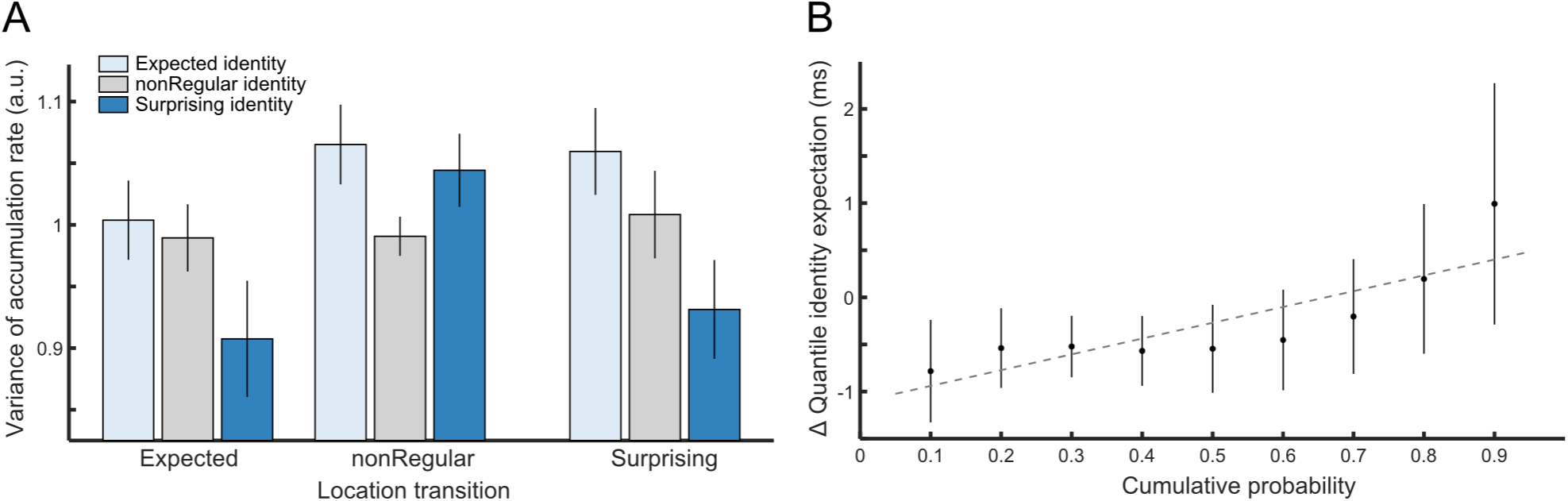
**Panel A.** Mean variance of accumulation rate in the nine cells from experimental conditions. Variance was higher for expected-identity than surprising-identity trials producing a main effect of Identity (see text). **Panel B.** Difference in SL quantiles between trials with expected and surprising identity. The difference increases from negative to positive values, indicating SL values were more dispersed when the identity transitions were expected.

Taken together with the fact that accumulation rate did not differ between expected and surprising trials, the difference in σ_*µ*_ points to an increased variability of SL values. This can be graphically visualized by the delta-plot (Schwarz & Miller, 2012) presented in Figure 6B, which presents the difference between expected- and surprising-identity condition across quantiles of the saccade latency distributions. The delta-plot shows that the difference between SL for expected and surprising trials changed from negative values for the lower quantiles (where expected-identity produced faster responses) to positive values for the upper quantiles (where expected-identity produced slower responses). The correlation between the *expected identity*− contrast and quantile was statistically significant, *Spearman’s rank* = 0.87, *p* = 0.005. This suggests that participants were engaged in a cognitive process that produced a larger set of outcomes for expected trials than for surprising trials.

#### Variance of Decision threshold

We did not find any impact of Location or Identity on the variance of decision threshold (3 × 3 ANOVA, all *F*_*s*_ < 1, *p*_*s*_ > .1). This suggests that our experimental manipulations impact exclusively the evidence accumulation stage.

### Anticipatory fixation offsets track location and identity regularities

Anticipatory fixation offsets are small gaze deviations from the fixation center that have been shown to track the next target location, prior to its appearance (Notaro et al., 2019). A preliminary analysis (see *Appendix*) showed that in this study, these offsets were not aligned with the location of the most probably future target. Instead, gaze offsets in the *x*-direction shifted towards the screen side opposite to that of the last presented target. This could reflect the fact that in both the regular and non-regular conditions, the mean probability of having the next target presented on the alternate side was 66%.

We examined whether gaze offsets could present a signature indicative of tracking location regularity, and whether identity regularity impacted gaze offsets as well.

#### Steady-state analysis

To determine whether gaze tracked the location transition structure of the experimental series, we first constructed series consisting of one gaze offset measure per trial. We then derived the power spectral densities of these series in the *x* − *axis* (see *Methods*). This evaluates whether there is a recurrence frequency that shows particularly high power.

We focused our analysis on the power density at the frequency *f*_*tag*_ = 1*/*3(*trials*^−1^). As can be seen in the power spectra derived from the target-locations themselves (see Appendix and Appx.1), this is the frequency in which there is a peak when location is regular. As shown in Figure 7A, the power spectra of the measured gaze-offset time series also peaked at *f*_*tag*_, with apparently little impact of whether identity was regular or not. This suggests that anticipatory gaze locations tracked the recurrence cycles of the Markov process that generated the targets’ locations.

**Figure 7.**
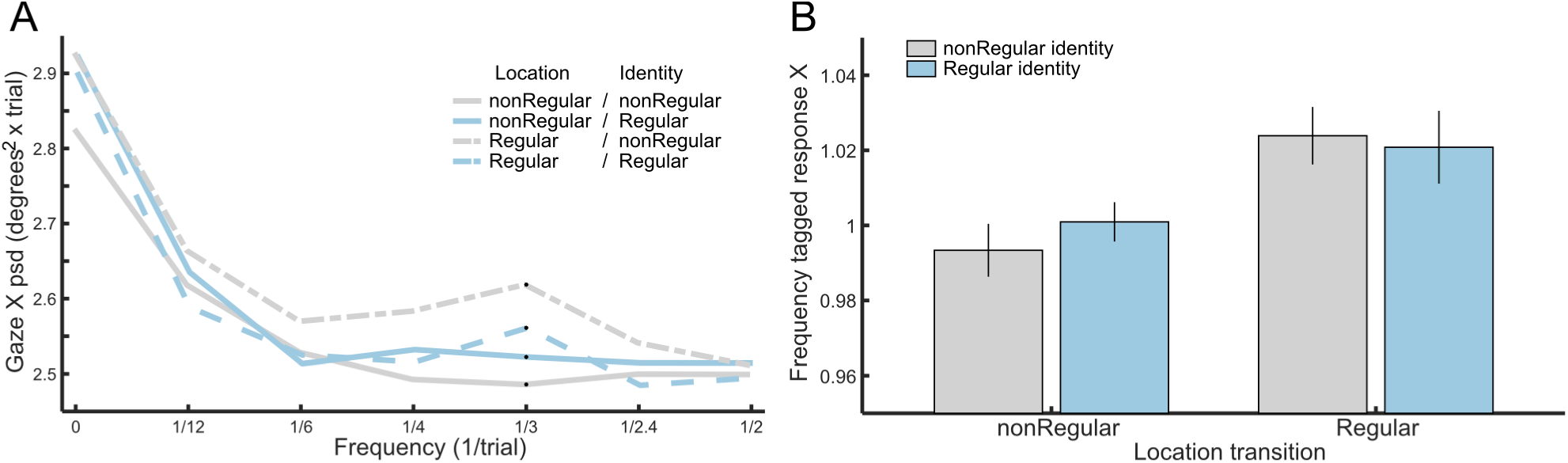
Results of steady-state analysis of gaze offsets. **Panel A.** Mean power-spectral density (PDS) plots of anticipatory gaze *X* positions in the four conditions with regular and nonRegular target locations (respectively dotted and continuous lines) and with regular and nonRegular identities (respectively blue and grey lines). The black circles indicate the power densities at the tagged frequency *f*_*tag*_ = 1*/*3(*trials*^−1^). **Panel B.** Mean frequency tagged responses (FTR) of anticipatory gaze X in the four conditions (blue and grey bars respectively indicate regular and nonRegular identity). The Frequency Tagged Responses is defined here as the ratio between the power at *f*_*tag*_ and the mean power in the closest four frequencies.

To evaluate these patterns statistically, we considered the frequency tagged response (*FTR*_*x*_ at *f*_*tag*_ = 1*/*3(*trials*^−1^), see *Methods*), using a two-way ANOVA with two factors (Location and Identity) each with two levels (nonRegular, Regular). The data analyzed are presented in Figure 7B. We found that only Location was significant, *F*(1, 39) = 12.16, *p* < .001. A similar − finding – a main effect of Location – was obtained by analyzing the series of gaze offsets in the *y axis*: *FTR*_*y*_, *F*(1, 39) = 12.93, *p* < .001.

#### Analysis of variance of AFO values

As mentioned above, an omnibus ANOVA indicated that gaze offsets in the *x*-direction tended to track the screen-side opposed to the last target. We quantified this, by coding as positive AFO the gaze offset in the direction of the previous stimulus, negative otherwise (Notaro et al., 2019). In all the four experimental conditions, AFOs were significantly negative: when location and identity were not regular, *M* = −0.150 ±.026°, *t*(39) = 5.78, *p* < .001, *d* = 0.91; when only location was regular, *M* = −0.130 ±.024°, *t*(39) = 5.40, *p* < .001, *d* = 0.85; when only identity was regular, *M* = −0.159 ±.026°, *t*(39) = 6.05, *p* < .001, *d* = 0.96; and when both location and identity were regular, *M* = − 0.130 .026°, *t*(39) = 4.95, *p* < .001, *d* = 0.78.

A 2 × 2 ANOVA with two factors (Location and Identity) of two levels (not regular, regular) identified only a significant main effect of Identity, *F*(1, 39) = 5.57, *p* = .020. The Location factor and the interaction term were not significant (*F*_*s*_ < 1). This demonstrates that identity regularities modulated anticipatory gaze location prior to target appearance.

## Discussion

Orienting our eyes to changes in our surrounding environments requires substantially longer time than needed to implement a saccade to the intended target. It is thought (Noorani & Carpenter, 2016) that saccading to a target of interest exceeds the time needed for their physical execution by tens of milliseconds, and that this delay is associated with what is fundamentally a decision-related process that various external or internal factors can bias. One of the main factors that impacts stimulus-guided oculomotor reactions and responses in general, are predictions based on prior knowledge (Friston, Kilner, & Harrison, 2006).

It is well established that knowledge about the location of a future target speeds up target-related responses, either when that knowledge is acquired via learning or when it is communicated on a trial-by-trial level via diagnostic cues. However, while many studies have studied how stimulus processing is impacted by the ability to predict a single stimulus dimension such as spatial location or identity, the functional and neurobiological organization that supports learning of regularities in more than one dimension, as well as multi-dimensional predictions, is poorly understood. To investigate this, we employed a probabilistic design in which participants were presented with visual series that manifested probabilistic regularities about future targets’ location and/or identity (as in Davis & Hasson, 2018). Participants were only asked to saccade to the single target (presented without any distractors), which only required attention to the target location.

We concentrated on saccade latencies (SL) because SL reflect the earliest stimulus-triggered behavioral responses. Applying ELATER (Nakahara et al., 2006), a simple decision model that quantifies three components that contribute to saccade latencies, produced new understandings of how knowledge of location and identity is integrated in the earliest stages of stimulus-guided responses. We found that learning the probabilistic transition structure of location and identity translates into interactive impacts of location and identity predictability. Our findings further suggest that some of these interactive effects occur because learning identity-regularities in and of themselves produce a cognitive load that impacts SL, as seen in the fact that identity regularities negatively impacted saccades to surprising locations, and also impacted on fixation offsets before the appearance of the target. Our results also replicated our recent work (Notaro et al., 2019) showing that anticipatory predictions of target locations can be read-off from the eye position during fixation, prior to target presentation. In what follows we describe how the findings expand on conclusions of prior work and discuss the theoretical implications.

### Validation and extension: regularities in location stream produce faster responses to targets appearing in expected locations and anticipatory fixation offsets

Saccade latencies tracked the location-transitions. Latencies were fastest to expected trials in regular series (lowest surprise), slowest to unexpected trials in regular series (highest surprise), and demonstrated intermediate values for trials in cases where location was non-regular and could not be predicted (mid-level of surprise). Accumulation rates quantified via ELATER closely tracked the SL findings for location-regularity.

Notaro et al. (2019) have shown that visual series that contain statistical regularities about target location produce Anticipatory Fixation Offsets (AFOs) during fixation, prior to target presentation, in the direction of the expected target location. In that study transition constraints held between two screen sides, whereas in the current study they held between four specific locations. To evaluate if the experimentally-induced transition structure modulated AFOs, we implemented a ‘steady-state’ analysis that quantified recurrence patterns in the gaze positions. We documented regularity-tracking anticipatory fixations, evident in a stronger power at the Markov recurrence frequency (modal value: 3 trials) when location was regular as compared to when it was not. This result corroborates Notaro et al.’s in demonstrating that anticipatory gaze patterns, in absence of any stimulus-guided response, can contain information about learning the input’s transition structure. These results, taken together, show that participants attended the series and followed instructions.

### Identity-regularity impacts saccade latencies via specific and non-specific routes

Because we examined responses to predictable and less predictable stimuli in the context of learning we could differentiate between two routes through which series-regularity could impact behavior: specific and non-specific effects.

#### Differentiating specific and non-specific routes for regularity effects

*Specific Effects* of regularity differentiated expected (high-predictable) transitions from surprising (low-predictability) transitions in series where a dimension (here, location or identity) was predictable. From a functional perspective, these effects may reflect evaluation of stimulus features against an anticipatory prediction, or processes related to generation and propagation of error/surprisal terms.

*Non-specific Effects* of regularity, on the other hand, differentiated non-predictable events (in random series) from both expected and surprising events in regular series, but without differentiating the latter two. To our knowledge, these effects have rarely been identified in the context of statistical learning. Formally, they correlate with learnability (i.e., the complexity of the specific set of constraints that needs to be learned), input-entropy, regularity or any other formal property that differentiates inputs that are more statistically regular from inputs that are less regular. From a functional perspective, non-specific effects could reflect learning and updating processes triggered by regular series independent of the surprisal of each input. They could also reflect attentional or memory processes that are more strongly expressed when input series contain strong associations. For instance, Zhao, Al-Aidroos, and Turk-Browne (2013) demonstrated that attention is drawn to regular, task-irrelevant shape streams, and Otsuka and Saiki (2016) have shown that items presented in regular series are better remembered than those presented within random series.

Separating specific from non-specific effects is important because they could load on different computations, and likely involve different neurobiological systems. Non-specific effects can be considered a minimal signature of learning, because there is no indication that learning has been translated into the sort of adaptive reactive behavior that improves stimulus-processing. In contrast, specific effects suggest that individuals are using their knowledge in a way that impacts trial-level assessment and in this way differentiates between expected and surprising events.

#### Non-specific effects of identity-regularity

Several indicators showed that participants were sensitive to the transition structure that existed between the identities of the stimuli (here, images drawn from four categories). In the analysis of accumulation-rate, we found a non-specific effect of Identity regularity: there was a main effect of Identity which did not track surprisal, and it was further modulated by a significant interaction between Location and Identity. The interaction was produced because the existence of identity-regularities strongly impacted saccades to surprising locations, but had no impact when targets were presented at expected or non-regular (non-predictable) locations.

Because saccades to surprising locations were associated with the slowest latencies, we investigated whether the determining factor was location-surprisal or the time to execute the saccade. We performed this analysis because prior eye-tracking studies have shown that top-down modulations such as implementing top-down control, integration of stimulus-value information, or impact of semantic priming are more strongly reflected in slower than faster saccades (e.g., Van Zoest & Donk, 2008; Schütz et al., 2012; Hoedemaker & Gordon, 2017). We performed median splits on saccade-latency distributions for saccades to surprising, expected, and non-regular locations. While our findings do not rule out a time-to-execute account, they do support the idea that this was the determining factor. When examining saccades to expected or surprising locations there was no statistical difference between identity costs for the slower and faster splits (though for both, identity costs were numerically greater for the slower splits). When examining saccades to non-regular locations, we found that identity-regularity appeared to produce a penalty for the slower split, but (a significant) facilitation for the slower split. This pattern was different from that found for the expected- and surprising-locations conditions. In addition, when examining cross-participant correlations between the magnitude of identity-costs across conditions, we found significant correlations between the costs associated with saccades to surprising and expected locations, but neither condition correlated with costs found in the non-regular condition. In all, this suggests that regularities in the identity stream impacted saccades to non-regular locations in a different way than they impacted saccades to expected or surprising locations.

We also found that the pattern of anticipatory fixations was impacted by identity-regularity. Specifically, identity-regularity reduced the bias to move the anticipatory fixation to the side opposite of that of the prior target.^6^ This is a clear sign that regularities in the identity stream impact stimulus-independent processes.

These findings are consistent with neurobiological data reported in Davis and Hasson (2018). That study collected fMRI data in a paradigm with a similar design to that used here, with the exception that targets were viewed passively and foveally, without initiating saccades. That study showed that when compared to series where neither Location or Identity was predictable, both the predictable-location and predictable-identity series produced reduced activity in multiple, partially overlapping brain regions. The dual-predictability condition – in which both location and identity were predictable – produced more modest signatures of reduced activity. Furthermore, in several brain areas (including frontal, inferior parietal and occipital regions) dual-predictability produced greater activity than the two conditions where either location or identity were predictable. This was interpreted as signaling a bottleneck in processing, associated with managing learning and prediction in two separate dimensions of the input stream.

Those findings are completely consistent with the current report, where we find that processing predictable streams of identity-information produced different effects depending on whether the target location was not-predictable or conversely, expected or surprising. This was most strongly seen in the current study in the negative impact of identity-regularity on action planning when the location prediction is disconfirmed. The anterior cingulate cortex may be involved in this bottleneck as it is related to both oculomotor control (Paus, Petrides, Evans, & Meyer, 1993) and surprisal detection (Hayden, Heilbronner, Pearson, & Platt, 2011), and has functional connections with inferior temporal cortex areas (Van Hoesen, Morecraft, & Vogt, 1993) sensitive to object categories (Haxby, Gobbini, Ishai, Schouten, & Pietrini, 2001).

#### Specific effects of identity regularity

In addition to the non-specific effects of identity-regularity summarized above we documented a specific effect of identity-regularity. We found that expected-identity transitions were associated with greater variance of accumulation-rate values than surprising-identity transitions. We interpret this finding as indicating that expected and surprising identity transitions produced similar sorts of processing, but that the expected condition produced additional types of cognitive outcomes, leading to greater variance. One possibility, which should be evaluated in future work, is that these effects are linked to an internal signal that scales with the match between the presented target and the predicted target. In the expected condition, the match could subsume a relatively large range of values, ranging from very strong to very weak matches (an analog in memory research would be the familiarity signal), whereas in the surprising condition, the match would subsume a lower range of values.^7^

This interpretation would be consistent with decision-making research about memory contents. That work (Starns, Ratcliff, & McKoon, 2012) found that when performing old/new memory judgments during a memory test, the variance of response latencies to old items strongly exceeds the variance associated with new items. This suggests that the variance of the match between the probe (the presented item) and the memory trace (in our case, the content of the prediction) is greater for Old (expected) than New (surprising) items.

The fact that we find a differentiation of expected vs. surprising-identity on saccade latencies is consistent with prior empirical data suggesting that category identity of pictorial stimuli can be very rapidly extracted (Grill-Spector & Kanwisher, 2005; Bar, 2004), and certainly within the time frames identified here. Behavioral measures using saccade-to-target show that categorization and saccade execution to correct categories (presented parafovealy at 6 degrees) can occur within ∼ 120 ms, though modal values are ∼ 150ms (Kirchner & Thorpe, 2006). These values are in the range of the saccade latencies we document here which were around 144ms (for the fastest condition; where locations were expected). The idea that familiarity can underlie differences in variance of accumulation rate at these early time points should addressed in the future, but related EEG work has suggested that expectations about object identity impact activity in category-specific areas at ∼170ms post stimuli (Aranda, Madrid, Tudela, & Ruz, 2010). Our findings regarding the specific effect of identity-regularity are somewhat independent of the literature on whether the coding of object location and object identity is performed serially or in parallel (Ullman, 2007; Schneider, 1995; Walther & Koch, 2006), because our study speaks to how early current object identity is evaluated against a prior template (Bar, 2003), perhaps during saccade planning. This evaluation could be mediated by the orbito-frontal cortex, which has been shown to rapidly integrate information at low spatial frequencies (Kirchner & Thorpe, 2006; Bar et al., 2005).

### Relation to prior studies examining joint processing of location and identity

As already reviewed in the Introduction, several studies have examined potential interactions between learning spatial and identity sequences (e.g., Mayr, 1996; Deroost & Soetens, 2006; Remillard, 2017), and other studies documented how individuals respond to targets after being provided cues that contain information about the target’s location and/or identity (e.g., Mattler, 2003; Bruhn & Bundesen, 2012; Egner et al., 2008). Prior studies have two main limitations in relation to understanding learning and the generation of multimodal predictions: *i*) all studies used stimulus-response paradigms that, beyond perceptual processing of item features, necessitate a Stimulus-Response mapping; that is, assigning a stimulus to a category and then selecting the proper response from a set of responses, and *ii*) cue-target paradigms in particular sample behavioral responses to exogenous cues, independently of learning. The study by Remillard (2017) – which was the most similar to our study in orthogonally manipulating spatial and identity probabilities – necessitated identity judgments.

These differences aside, our results can be viewed as being consistent with the original findings reported by Mayr (1996), who found no indication that learning identity sequences interferes with learning of spatial sequences. That study used orthogonal location/identity series to deterministically assign location/identity transitions. When location sequences are deterministic, any subsequent spatial transition is by definition a high probability one. As shown in the current study, the interaction between location and identity information was only found when saccading to surprising locations, in which case regularity in identity series produced even slower responses. Thus, prior conclusions about non-interactivity could be attributed to the use deterministic series that contained only high-probability (certain) transitions, and that did not produce a processing bottleneck.

### Limitations

Our study shows how identity regularities interact with the saccade generation process. Departing from previous studies, we chose to not include any decision about the image identity, but future studies could modify the paradigm to draw participants’ attention toward identity, and in this way probe for the effect of Identity in a different manner. The experimental paradigm could be modified to overcome some current limitations. For example, the gap time between fixation and target-presentation can be varied between trials. This could produce a greater variance of saccade latencies and in this way inrease the sensitivity of some statistical analyses. It would also be useful to understand how the experimental manipulations impacted dwell-times on the target-images once they were fixated. In the current study, the pace of target presentation was rapid and oftentimes the image disappeared before participants could initiate a saccade away from the image. This reduced the validity of such an analysis for purposes of studying the impact of series-regularity on stimulus-processing.

### Summary

We examined whether statistical regularities in object-location and item-identity produced independent or interactive processes. We used a saccade-to-target procedure and we modeled saccade latencies as decisions about target location (ELATER). Regularities in the identity stream decreased the rate of evidence accumulation when the target location was surprising. Additionally, expected (vs. not expected) identities increased the variance of accumulation rate, indicating a larger range of outcomes in the underlying decision process. Joint impacts of location-regularity and identity-regularity were also identified in analysis of anticipatory fixation offsets. In all, our findings demonstrate a strong interaction between the processing of regularities in object location and identity during stimulus-guided saccades, and suggest these regularities also impact anticipatory, non-stimulus-guided fixations.

### Appendix

#### Sequences generation and image preparation

To equally represent all the possible eye trajectories, in the regular condition we generated series of target locations using four different transition matrices. Specifically, in addition to the matrix *M*_*Reg*1_ presented in the main text, we used also *M*_*Reg*2_ below, and two transpositions of those matrices, 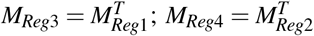.

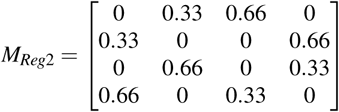

Note that since each element *a*_*i j*_ of a transition matrix is equal to the element *b* _*ji*_ of its transpose, we generated location sequences where two eye trajectories with opposite directions (i.e. 1-2 and 2-1) had the same probability to occur. When considered across these four matrices, the proportion of pair-wise transitions was equal for the Regular and nonRegular processes. The nonRegular matrix (in the main text) is in fact the average of the the four Regular matrices. This meant that statistical structure was not confounded with the pair-wise movements across trials. The 120-elements sequences were generated joining three 40-elements sequences with the intended statistical features (marginals and transition probabilities). To make sure that the location and identity series were not mutually informative, in each experimental condition, a sequence of target locations was paired with the least correlated sequence of target identity among a set of hundreds of sequences generated with a given transition matrix.

To avoid an edge contrast between the images and the display background (set to mid gray), we isolated the foreground with a custom software and we then added as background the same uniform mid gray of the display. To equalize image statistics we set the foregrounds to have the same mean luminance (125/255) and standard deviation (70/255) changing the third coordinate to HSV representation, and then transforming back to RGB. For each image we calculated the mean pixel position weighted with the luminance and we used these points to locate the images in the display. Images were resized to have the same size and pixels resolution (450 *×* 600) and approximately the same number of pixels in the foreground.

#### Temporal characterization of location series

A property of Markov processes is that processes with different levels of transition probabilities have different periods of repetitions, which translates into different autocorrelation functions (and analogously, power spectra). Given a transition matrix *P*, its element *p*_*i j*_ is the probability of the target appearing at the location *l* _*j*_ if on the current trial it is in the location *l*_*i*_. Similarly the element *p*_*i j*_ of the matrix *P*^*n*^ = *PxP*^*n*−1^ indicates the probability of the target appearing after *n* transitions at the location *l* _*j*_ if it is in the location *l*_*i*_. Consequently, for the considered *Reg* matrices, the probability to complete the most likely screen side transition after *n* trials, has a maximum at n=1 (p=0.66), but peaks after three more transitions (n=4, p=0.39) (Figure Appx.1B, grey circles). Conversely, for the *nonReg* matrix, the probability to complete an allowed transition after *n* transitions has a peak after two trials (n=3, p=0.26) (Figure Appx.1B, black circles). This recurrence characteristic of Markov processes underlies the temporal features of any possible allowed series of transitions.

**Figure Appx.1.**
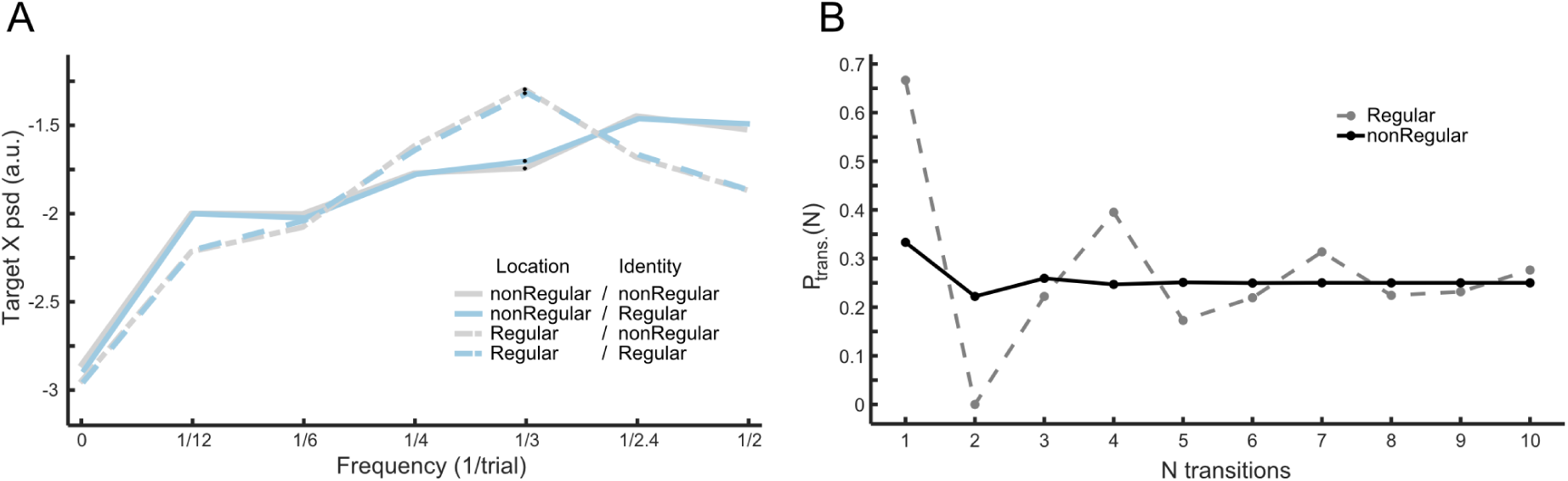
**A.** PSD of X-axis target location **B.** Temporal characterization of Markov processes.

#### Gaze offsets does not directly signal screen side of most likely subsequent target

In the condition with Regular location transitions, on each trial there was a probability of 66% of transitioning to one location, 33% probability of transitioning to another, and 0% transition to a third (in addition, repeats were never allowed). To understand if anticipatory gaze offsets were aligned to the location of the most probably future target, we partitioned all the trials into 4 bins – right/left × top/bottom – depending on the most probable target location in the next trial. If participants’ gaze tracked the most likely future position then on first approximation, gaze location should show greater bias towards the right side when the *P* = 66% transition is expected to be on the right than when the *P* = 66% transition is expected to ben the left, and similarly, show greater bias towards the top/bottom of the screen depending of the expected vertical position of the next most probable target. An Analysis of Variance with three factors (horizontal position of next most probable target, vertical position of next most probable target, and screen side of just presented image) did not indicate that anticipatory gaze was biased towards the horizontal position of the next most likely target, since the first factor and its interaction were not significant (*F* < 1). The only significant effect was the screen side of previous screen *F*(1, 39) = 80.4 *p* < 10^−6^, because there was a general shift towards the side opposite to that of the last presented target.

### Graphical depiction of ELATER parameters

**Figure Appx.2.**
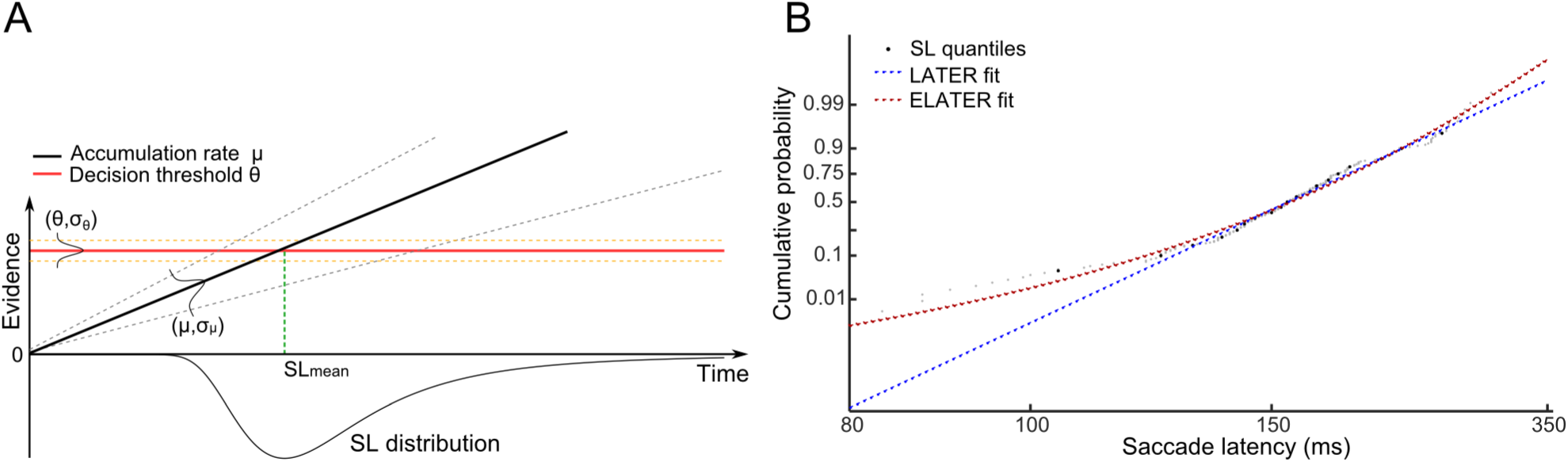
ELATER model. **Panel A:** During each trial, starting from t=0 (visual target presentation) evidence (y-axis) is linearly accumulated (black line) with a constant rate extracted from a normal distribution with average *µ* and standard deviation σ_*µ*_. A decision is taken when evidence reaches the decision threshold *θ* (red line), from a normal distribution with mean equal to one and standard deviation σ_*θ*_. **Panel B**: reci-probit plot of SL data (grey dots) fit with LATER model (straight blue line) and ELATER model (curved red line). ELATER fit is calculated in quantiles of probabilities spaced at 0.05 (black dots).

## Notes

1 In this study valid cues provided absolutely certain foreknowledge, whereas invalid cues were completely uninformative

2 We did not consider the angle value because it provides less information about the strength of the anticipatory pattern (a single angle is consistent with multiple {*X,Y*} tuples), and because minor changes in *X* or *Y* would translate into large angular differences due to the relatively small deviations from center.

3 The parameter values obtained this way highly correlated (all *rho* > .4, all *p* < .01) with the mean of the parameters obtained by separately applying ELATER to data with expected and surprising identity.

4 Similar findings were identified when using Saccade Latencies to identify identity costs. When SL, the correlation between *idcost*_exp_ and *idcost*_surp_ was− 0.43(*p* = .0052), and the correlation between *idcost*_nonReg_ and *idcost*_surp_ was 0.21(*p* > .1)

5 Note that averages of saccade latencies were used because it was not possible apply the ELATER decision model to median-split data

6 This bias was licensed by the fact that, on average, in all conditions there was a greater probability of a target presented on the other side than the same side as the prior target

7 We note that the Expected condition consisted of more trials than the Surprising condition (by definition) and for this reason, all else being equal (i.e., if sampling from the same sampling distribution of saccade latencies), the Surprising condition should have been associated with greater variance due to the lower number of trials.

